## Supplementary tables for "Bidirectional alterations in brain temperature profoundly modulate spatiotemporal neurovascular responses in-vivo: Implications for theragnostics"

Supplementary Table 1: 2s Baseline and response values

| | 44°C | 40 °C | 37 °C | Ambient | 20 °C | 10 °C | 6 °C | Single Factor Anova Result |
|---|---|---|---|---|---|---|---|---|
| Actual brain temperature (°C) | 39.8 ± 0.2 | 37.5 ± 0.2 | 36.0 ± 0.2 | 28.4 ± 0.8 | 23.4 ± 0.3 | 15.3 ± 0.5 | 12.5 ± 0.6 | F=551.1 df=6 $p=6.4x10^{-27}$ |
| Change in brain temperature (oC) | -0.019 ± 0.004 | -0.008 ± 0.003 | 0.008 ± 0.002 | 0.051 ± 0.007 | 0.096 ± 0.020 | 0.054 ± 0.019 | 0.013 ± 0.004 | F=9.93 df=6 $p=2.9x10^{-6}$ |
| Brain tissue $O_2$ measurement (mmHg) | 37.6 ± 3.0 | 33.6 ± 4.0 | 30.6 ± 3.9 | 12.4 ± 3.1 | 15.1 ± 2.4 | 9.1 ± 1.2 | 8.2 ± 1.0 | F=13.69 df=6 $p=4.3x10^{-7}$ |
| Change in Tissue oxygen (Fractional) | 1.021 ± 0.007 | 1.036 ± 0.008 | 1.052 ± 0.024 | 1.046 ± 0.017 | 1.029 ± 0.013 | 0.977 ± 0.010 | 0.991 ± 0.004 | F=3.51 df=6 p=0.009 |
| Hbt Onset (s) | 0.54 ± 0.14 | 0.64 ± 0.07 | 0.45 ± 0.05 | 0.79 ± 0.13 | 1.81 ± 0.13 | 3.40 ± 0.74 | 4.82 ± 1.77 | F=4.15 df=6 p=0.003 |
| Hbt Time to peak (s) | 2.59 ± 0.05 | 2.79 ± 0.22 | 2.92 ± 0.19 | 3.83 ± 0.49 | 5.52 ± 0.27 | 9.96 ± 1.84 | 13.46 ± 6.76 | F=11.52 df=6 $p=6.3x10^{-7}$ |
| Hbt Peak magnitude (Fractional) | 1.028 ± 0.006 | 1.040 ± 0.006 | 1.042 ± 0.008 | 1.051 ± 0.013 | 1.056 ± 0.011 | 1.01 ± 0.002 | 1.003 ± 0.001 | F=5.48 df=6 p=0.0005 |
| Hbr dip magnitude (Fractional) | 1.009 ± 0.002 | 1.008 ± 0.002 | 1.005 ± 0.001 | 1.007 ± 0.001 | 1.014 ± 0.001 | 1.019 ± 0.003 | 1.011 ± 0.001 | F=7.01 df=6 $p=7.2x10^{-5}$ |
| LFP Magnitude (Volts) | 0.47 ± 0.04 | 0.52 ± 0.03 | 0.56 ± 0.01 | 0.54 ± 0.03 | 0.69 ± 0.04 | 0.64 ± 0.06 | 0.48 ± 0.05 | F=3.29 df=6 p=0.015 |
| LFP Minima (Volts) | -0.0017 ± 0.0003 | -0.0022 ± 0.0002 | -0.0025 ± 0.0003 | -0.0020 ± 0.0002 | -0.0019 ± 0.0002 | -0.0011 ± 0.0002 | -0.0006 ± 0.0001 | F=5.65 df=6 p=0.00066 |
| MUA Magnitude (Spikes total during impulse) | 156.46 ± 5.35 | 187.77 ± 5.5 | 187.74 ± 5.63 | 230.72 ± 7.11 | 243.52 ± 10.24 | 164.31 ± 16.58 | 61.62 ± 8.94 | F=33.38 df=6 $p=1.17x10^{-12}$ |

Supplementary Table 2: 16 Baseline and response values

| | 44 °C | 40 °C | 37 °C | Ambient | 20 °C | 10 °C | 6 °C | Single Factor Anova Results |
|---|---|---|---|---|---|---|---|---|
| Actual brain temperature (°C) | 39.4 ± 0.3 | 37.5 ± 0.2 | 36.0 ± 0.2 | 28.9 ± 0.7 | 23.4 ± 0.2 | 15.1 ± 0.4 | 12.5 ± 0.5 | F=652.7 df=6 $p=2.1x10^{-34}$ |
| Change in brain temperature (oC) | -0.01 ± 0.003 | -0.003 ± 0.002 | 0.015 ± 0.006 | 0.146 ± 0.016 | 0.252 ± 0.022 | 0.086 ± 0.021 | 0.076 ± 0.018 | F=33.45 df=6 $p=4.25x10^{-13}$ |
| Brain tissue $O_2$ measurement (mmHg) | 38.3 ± 4.1 | 35.3 ± 3.9 | 30.9 ± 3.6 | 14.0 ± 3.0 | 13.2 ± 2.1 | 8.2 ± 1.2 | 8.1 ± 1.1 | F=16.447 df=6 $p=6.7x10^{-9}$ |
| Change in Tissue oxygen (Fractional) | 1.068 ± 0.025 | 1.119 ± 0.038 | 1.1153 ± 0.053 | 1.175 ± 0.093 | 1.081 ± 0.038 | 0.980 ± 0.033 | 0.967 ± 0.014 | F=2.31 df=6 p=0.054 |
| Hbt Onset (s) | 0.47 ± 0.22 | 0.47 ± 0.09 | 0.56 ± 0.05 | 0.87 ± 0.20 | 1.86 ± 0.14 | 4.66 ± 0.31 | 6.51 ± 0.41 | F=89.24 df=6 $p=9.8x10^{-20}$ |
| Hbt Time to peak (s) | 3.07 ± 0.14 | 3.27 ± 0.12 | 3.77 ± 0.15 | 9.71 ± 1.89 | 10.13 ± 1.40 | 17.48 ± 1.36 | 23.79 ± 3.42 | F=19.21 df=6 $p=9.0x10^{-10}$ |
| Hbt Peak magnitude (Fractional) | 1.031 ± 0.005 | 1.046 ± 0.007 | 1.050 ± 0.009 | 1.079 ± 0.013 | 1.096 ± 0.014 | 1.020 ± 0.002 | 1.003 ± 0.001 | F=8.959 df=6 $p=6x10^{-6}$ |
| Hbr dip magnitude (Fractional) | 1.011 ± 0.003 | 1.004 ± 0.001 | 1.008 ± 0.001 | 1.009 ± 0.001 | 1.013 ± 0.002 | 1.024 ± 0.002 | 1.022 ± 0.002 | F=11.63 df=6 $p=4.0x10^{-7}$ |
| LFP Magnitude (Volts) | 0.27 ± 0.03 | 0.31 ± 0.03 | 0.34 ± 0.03 | 0.42 ± 0.02 | 0.47 ± 0.04 | 0.16 ± 0.03 | 0.087 ± 0.01 | F=21.01 df=6 $p=2.77x10^{-10}$ |
| LFP Minima (Volts) | -0.0014 ± 0.0002 | -0.0019 ± 0.0003 | -0.0020 ± 0.0003 | -0.0020 ± 0.0002 | -0.0016 ± 0.0002 | -0.0004 ± 0.0001 | -0.0002 ± 0.00005 | F=13.02 df=6 $p=1.09x10^{-7}$ |
| MUA Magnitude (Spikes total during impulse) | 152.80 ± 14.58 | 169.90 ± 6.97 | 173.93 ± 8.98 | 213.54 ± 8.58 | 216.59 ± 6.65 | 76.85 ± 15.93 | 31.09 ± 7.50 | F=43.81 df=6 $p=7.58x10^{-15}$ |

Supplementary Table 3: Significant post-hoc Tukey test results

| | Two Seconds (DF all 33) | Sixteen seconds(DF all 35) |
|---|---|---|
| Actual brain temperature | 44°C-37°C,p=0.0004<br>44°C-Amb, p<0.0001<br>44°C-20 °C, p<0.0001<br>44°C-10 °C, p<0.0001<br>44°C-6 °C, p<0.0001<br>40°C-Amb, p<0.0001<br>40°C-20 °C, p<0.0001<br>40°C-10 °C, p<0.0001<br>40°C-6 °C, p<0.0001<br>37°C-Amb, p<0.0001<br>37°C-20°C, p<0.0001<br>37°C-10°C, p<0.0001<br>37°C-6 °C, p<0.0001<br>Amb -20 °C, p<0.0001<br>Amb -10 °C, p<0.0001<br>Amb -6°C, p<0.0001<br>20°C-10 °C, p<0.0001<br>20°C-6°C, p<0.0001<br>10°C-6 °C, p=0.0033 | 44°C-40°C,p=0.0474<br>44°C-37°C, p<0.0001<br>44°C-Amb, p<0.0001<br>44°C-20 °C, p<0.0001<br>44°C-10 °C, p<0.0001<br>44°C-6 °C, p<0.0001<br>40°C-Amb, p<0.0001<br>40°C-20 °C, p<0.0001<br>40°C-10 °C, p<0.0001<br>40°C-6 °C, p<0.0001<br>37°C-Amb, p<0.0001<br>37°C-20°C, p<0.0001<br>37°C-10°C, p<0.0001<br>37°C-6 °C, p<0.0001<br>Amb -20 °C, p<0.0001<br>Amb -10 °C, p<0.0001<br>Amb -6°C, p<0.0001<br>20°C-10 °C, p<0.0001<br>20°C-6°C, p<0.0001<br>10°C-6 °C, p=0.0023 |
| Change in brain temperature | 44°C-Amb, p=0.0177<br>44°C-20 °C, p<0.0001<br>44°C-10 °C, p=0.0123<br>40°C-Amb, p=0.0316<br>40°C-20 °C, p<0.0001<br>40°C-10 °C, p=0.0212<br>37°C-20°C, p=0.0003<br>20°C-6°C, p=0.0008 | 44°C-Amb, p<0.0001<br>44°C-20 °C, p<0.0001<br>44°C-10 °C, p=0.0035<br>44°C-6 °C, p=0.011<br>40°C-Amb, p<0.0001<br>40°C-20 °C, p<0.0001<br>40°C-10 °C, p=0.0075<br>40°C-6 °C, p=0.0208<br>37°C-Amb, p<0.0001<br>37°C-20°C, p<0.0001<br>Amb -20 °C, p=0.0009<br>20°C-10 °C, p<0.0001<br>20°C-6°C, p<0.0001 |
| Actual Tissue oxygen | 44°C-Amb, p=0.0002<br>44°C-20 °C, p=0.0011<br>44°C-10 °C, p<0.0001<br>44°C-6 °C, p<0.0001<br>40°C-Amb, p=0.0006<br>40°C-20 °C, p=0.0032<br>40°C-10 °C, p<0.0001<br>40°C-6 °C, p<0.0001<br>37°C-Amb, p=0.0039<br>37°C-20°C, p=0.019<br>37°C-10°C, p=0.0005<br>37°C-6 °C, p=0.0002 | 44°C-Amb, p=0.0001<br>44°C-20 °C, p<0.0001<br>44°C-10 °C, p<0.0001<br>44°C-6 °C, p<0.0001<br>40°C-Amb, p=0.0009<br>40°C-20 °C, p=0.0005<br>40°C-10 °C, p<0.0001<br>40°C-6 °C, p<0.0001<br>37°C-Amb, p=0.0125<br>37°C-20°C, p=0.0078<br>37°C-10°C, p=0.0004<br>37°C-6 °C, p=0.0003 |

| | | |
|---|---|---|
| Change in Tissue oxygen | 37°C-10 °C, p=0.017<br>Amb -10 °C, p=0.037 | NO Sig post-hoc results |
| Hbt onset | 44 °C-6 °C, p=0.0352<br>40 °C-6 °C, p=0.0167<br>37 °C-6 °C, p=0.019<br>Amb-6°C, p=0.0229 | 44°C-20 °C, p=0.0078<br>44°C-10 °C, p<0.0001<br>44°C-6 °C, p<0.0001<br>40°C-20 °C, p=0.0077<br>40°C-10 °C, p<0.0001<br>40°C-6 °C, p<0.0001<br>37°C-20 °C, p=0.0149<br>37°C-10 °C, p<0.0001<br>37°C-6 °C, p<0.0001<br>Amb -10 °C, p<0.0001<br>Amb -6°C, p<0.0001<br>20°C-10 °C, p<0.0001<br>20°C-6°C, p<0.0001<br>10°C-6 °C, p=0.0002 |
| Hbt time to Peak | 44°C-10 °C, p=0.0095<br>44 °C-6 °C, p<0.0001<br>40°C-10 °C, p=0.0039<br>40 °C-6 °C, p<0.0001<br>37 °C-10 °C, p=0.0047<br>37 °C-6 °C, p<0.0001<br>Amb -10 °C, p=0.019<br>Amb -6°C, p<0.0001<br>20 °C-6 °C, p=0.011 | 44°C-10 °C, p<0.0001<br>44 °C-6 °C, p<0.0001<br>40 °C-10 °C, p<0.0001<br>40 °C- 6 °C, p<0.0001<br>37 °C-10 °C, p=0.0001<br>37 °C-6 °C, p<0.0001<br>Amb -6°C, p<0.0001<br>20 °C-6 °C, p=0.0001 |
| Hbt magnitude | Amb -6°C, p=0.031<br>20 °C-10 °C, p=0.0122<br>20 °C-6 °C, p=0.0024 | 44 °C -Amb, p=0.0076<br>44°C-20 °C, p= 0.0001<br>40 °C-Amb, p=0.0273<br>40°C-20 °C, p<0.0001<br>37°C-20 °C, p=0.0002<br>Amb -10°C, p=0.0256<br>Amb -6°C, p=0.0005<br>20 °C-10 °C,p<0.0001<br>20 °C-6 °C, p<0.0001 |
| Hbr Dip Magnitude | 44°C-10 °C, p=0.0181<br>40°C-10 °C, p=0.0017<br>37°C-20 °C, p=0.0196<br>37°C-10 °C, p<0.000<br>Amb -10°C, p=0.0008 | 44°C-10 °C, p=0.0015<br>44 °C-6 °C, p=0.01<br>40°C-10 °C, p<0.0001<br>40 °C -6 °C, p<0.0001<br>37°C-10 °C, p=0.0002<br>37°C -6 °C, p=0.0013<br>Amb -10°C, p=0.0005<br>Amb -6°C, p=0.0032<br>20 °C -10 °C,p=0.0123 |
| LFP Magnitude | 44°C-20 °C, p=0.0393<br>20 °C-6 °C, p=0.0213 | 44 °C–Amb, p=0.0146<br>44°C-20 °C, p=0.0007<br>44°C-6 °C, p=0.0017<br>40°C-20 °C, p=0.0078<br>40°C-10 °C, p=0174<br>40°C-6 °C, p=0.0001<br>37 °C -20 °C, p=0.0418 |

| | | |
|---|---|---|
| | | 37 °C -10 °C, p=0.0029<br>37 °C -6°C, p<0.0001<br>Amb -10 °C, p<0.0001<br>Amb -6°C, p<0.0001<br>20 °C -10 °C, p<0.0001<br>20 °C -6 °C, p<0.0001 |
| LFP Minima | 40°C-10 °C, p=0.0468<br>40 °C - 6 °C, p=0.0012<br>37 °C -10 °C, p=0.0055<br>37 °C -6°C, p=0.0001<br>Amb -6°C, p=0.0037<br>20 °C -6 °C, p=0.01 | 44°C-10 °C, p=0.03<br>44°C-6 °C, p=0.0042<br>40°C-10 °C, p=0.0004<br>40°C-6 °C, p<0.0001<br>37°C-10 °C, p=0.0001<br>37°C-6 °C, p<0.0001;<br>Amb -10 °C, p<0.0001<br>Amb -6°C, p<0.0001<br>20 °C -10 °C, p=0.0036<br>20 °C -6 °C, p=0.0004 |
| MUA Magnitude | 44 °C -Amb, p=0.0012<br>44°C-20 °C, p= 0.0001<br>44°C-6°C, p<0.0001<br>40°C-20 °C, p=0.009<br>40°C-6°C, p=<0.0001<br>37°C-20 °C, p=0.009<br>37°C-6°C, p=<0.0001<br>Amb -10 °C, p=0.0012<br>Amb -6°C, p<0.0001<br>20°C-10 °C, p<0.0001<br>20°C-6°C, p<0.0001<br>10°C-6°C, p<0.0001 | 44 °C -Amb, p=0.004<br>44°C-20 °C, p= 0.0023<br>44°C-10°C, p=0.0002<br>44°C-6°C, p<0.0001<br>40°C-20 °C, p= 0.0475<br>40°C-10°C, p<0.0001<br>40°C-6°C, p<0.0001<br>37°C-10°C, p<0.0001<br>37°C-6°C, p<0.0001<br>Amb-10°C, p<0.0001<br>Amb-6°C, p<0.0001<br>20°C-10°C, p<0.0001<br>20°C-6°C, p<0.0001 |
